# Neural lineage induction reveals multi-scale dynamics of 3D chromatin organization

**DOI:** 10.1101/004168

**Authors:** Aleksandra Pękowska, Bernd Klaus, Felix A. Klein, Simon Anders, Małgorzata Oleś, Lars M. Steinmetz, Paul Bertone, Wolfgang Huber

## Abstract

Regulation of gene expression underlies cell identity. Chromatin structure and gene activity are linked at multiple levels, via positioning of genomic loci to transcriptionally permissive or repressive environments and by connecting *cis*-regulatory elements such as promoters and enhancers. However, the genome-wide dynamics of these processes during cell differentiation has not been characterized. Using tethered chromatin conformation capture (TCC) sequencing we determined global three-dimensional chromatin structures in mouse embryonic stem (ES) and neural stem (NS) cell derivatives. We found that changes in the propensity of genomic regions to form inter-chromosomal contacts are pervasive in neural induction and are associated with the regulation of gene expression. Moreover, we found a pronounced contribution of euchromatic domains to the intra-chromosomal interaction network of pluripotent cells, indicating the existence of an ES cell-specific mode of chromatin organization. Mapping of promoter-enhancer interactions in pluripotent and differentiated cells revealed that spatial proximity without enhancer element activity is a common architectural feature in cells undergoing early developmental changes. Activity-independent formation of higher-order contacts between *cis*-regulatory elements, predominant at complex loci, may thus provide an additional layer of transcriptional control.

---

The three-dimensional organization of chromatin is associated with transcriptional regulation at multiple scales. At the level of whole chromosome territories, loci are positioned to transcriptionally permissive (euchromatic) or repressive (heterochromatic) environments (1–2). More locally at genomic scales up to ∼3 Mb, chromatin is organized into largely cell-type invariant topologically associating domains (TADs, 3–5) which are regions of preferential chromatin interactions. TADs comprise the majority of local *cis*-regulatory element associations (6–7). At single-gene resolution, *cis*-regulatory elements function in tandem by forming physical contacts, resulting in a locus-specific chromatin structure (1).

At each of these scales, chromatin structural features can undergo changes during cell differentiation. Known alterations include tethering of repressed loci to heterochromatin (8–10) or to the nuclear periphery (8–9, 11). Moreover, gene-specific studies have observed repositioning of individual loci with respect to the surface of chromosomes (12–14). Finally, at the level of interactions within TADs, *cis*-regulatory elements such as promoters and enhancers form activity-associated, cell type-specific contact networks (7, 15–16).

Previous studies have investigated the link between alterations of chromatin structure and transcriptional regulation during differentiation, either for targeted loci (17–19) or at a genome-wide scale (8). Recently, RNA polymerase II (Pol II)-mediated chromatin interactions were analyzed by the ChIA-PET technique in the context of cell fate specification (16, 20). These studies have increased our understanding of the involvement of cell type-specific and ubiquitous transcription factors in the establishment of chromatin structure (16–18). Moreover, use of ChIA-PET allowed the comparative analysis of active (bound by Pol II) enhancer-promoter pairs during differentiation (20) and in distinct cell types (16) and revealed highly dynamic repertoire of promoter-enhancer contacts, involved in the control of expression of cell type specific and broadly expressed genes. However, a comprehensive map of the relationships between 3D nuclear structure and cell fate decisions at different genomic scales is still lacking.

Here we employ genome-wide chromatin conformation capture coupled with high throughput sequencing to analyze the alterations of chromatin structure accompanying neural stem (NS) cell induction from mouse embryonic stem cells (ES). We show that dynamic repositioning of TADs to heterochromatic regions is rare and affects only a small fraction of TADs during differentiation. In sharp contrast, we observed widespread, activity-linked relocations of TADs to chromosomal surfaces during the ES to NS cell transition. At the level of chromosome territory architecture, comparative analysis of TAD associations reveals previously unappreciated modes of chromatin organization between pluripotent cells and multipotent progeny. We show that euchromatin more often forms strong contacts in ES cells. This observation may be related to the overall higher activity of euchromatin in pluripotent cells. Finally, by high-resolution analysis of the dynamics of promoter interactomes we discover frequent dissociation between enhancer element activity and formation of contacts with promoters. We show that this uncoupling between spatial proximity and *cis*-regulatory element activity is present at many developmentally regulated genes and is highly pronounced at complex, multi-enhancer loci.

## RESULTS

### Dynamic repositioning of TADs with respect to eu- and heterochromatin

We used Tethered Chromatin Conformation Capture (TCC, 21) to determine the chromatin structure of mouse embryonic stem (ES) cells and neural stem (NS) cells, which we derived directly from the ES cells by *in vitro* differentiation (**Fig. 1*A***, **Fig. S1**, SI Methods, (22–24). We generated TCC libraries for two biological replicates each of ES (46C, 22) and NS cells. After stringent quality control (**Fig. S2**), we analyzed ∼200 million sequence read pairs corresponding to putative interactions between individual HindIII restriction fragments (**Table S1**).

**Fig. 1.**
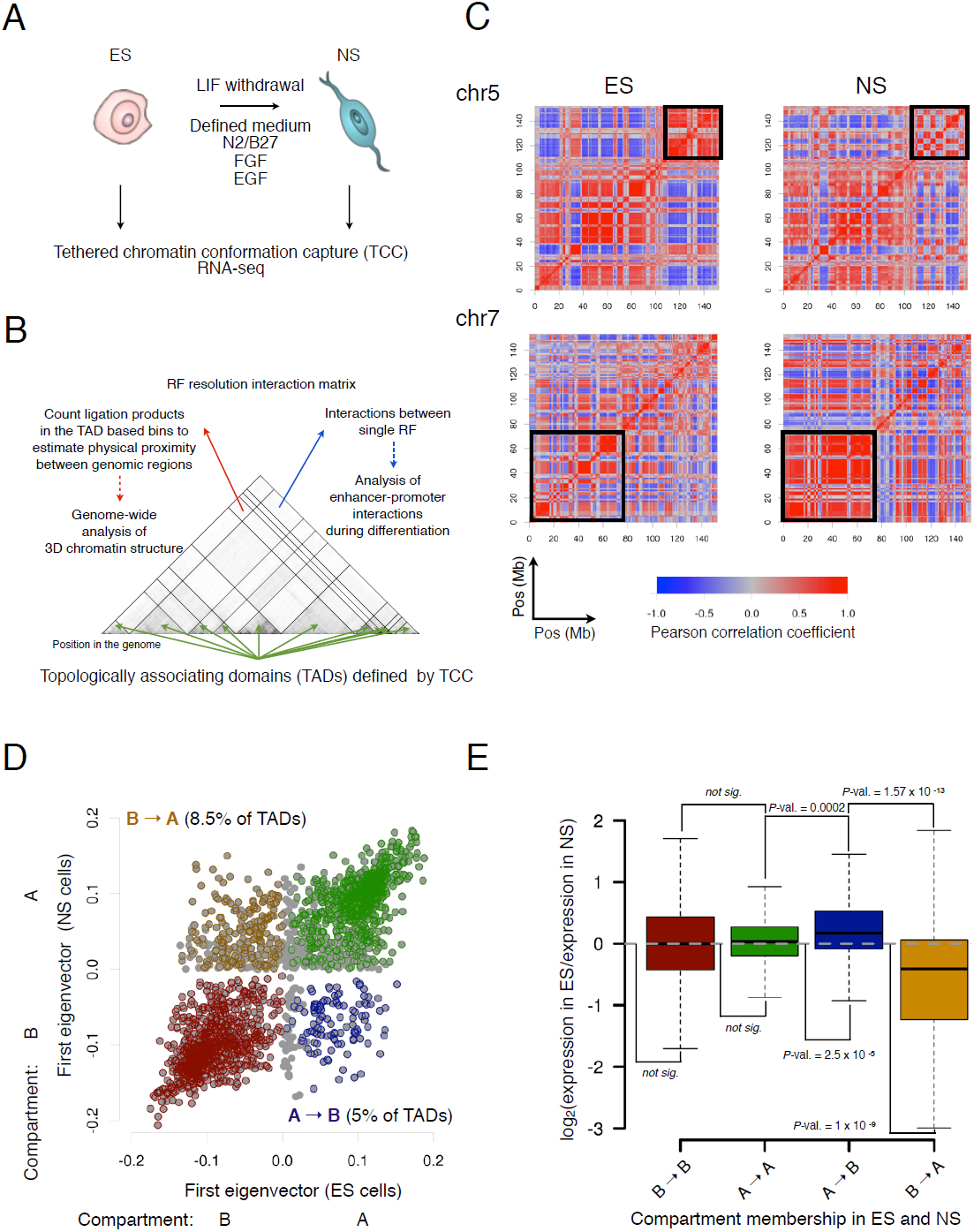
Shuttling of topologically associating domains (TADs) in and out of heterochromatic regions is associated with differentiation and with coordinate regulation of gene expression. (***A***) Design of the study. (***B***) Overview of the analyses. TAD boundaries were determined from the pooled dataset by a segmentation algorithm. The strength of interaction between each pair of TADs was measured by the count of ligation products in the corresponding rectangular bins. In parallel, physical interactions at restriction fragment resolution were analyzed to study promoter-enhancer interaction dynamics. (***C***) Correlation matrices of the normalized intra-chromosomal ligation frequencies at TAD resolution for chromosomes 5 and 7 reveal numerous zones of changes of interaction patterns (black squares). (***D***) Compartmental membership of TADs in ES and NS cells. Two compartments (A and B) were isolated from principal component analysis of the chromosome-wise correlation maps and RNA-seq data. The strength of association with the A and B compartment in both cell types is displayed. Encircled points indicate compartment assignment (Methods). (***E***) TAD-wise average gene expression changes between ES and NS cells. TADs were stratified by the compartment membership dynamics defined in panel ***D***. P-values: two-sided t-test.

To assess alterations in 3D chromatin structure accompanying the ES-to-NS cell transition, we inferred a partitioning of the genome into topologically associating domains (TAD, 3–5) at restriction fragment resolution separately for each cell population (**Fig. 1*B***, SI Methods, **Figs. S3 and S4**). In agreement with previous observations (3), TADs were largely cell-type invariant (**Fig. S5**). Therefore, to perform a direct comparative analysis of the TAD interactomes, we inferred a consensus set of 2232 TADs from the combined data. We used TAD coordinates as counting bins and determined the strength of pairwise TAD interactions based on the number of read pairs connecting them. The resulting interaction profiles were reproducible between replicates and consistent with published data (**Figs. S6 and S7**).

Association and dissociation of loci with heterochromatic regions has been linked to the regulation of gene expression in cell differentiation (8–10). Previous studies have reported the repositioning of developmentally downregulated genes to heterochromatin domains (10) or to the nuclear periphery (8–9, 11). However, because these efforts have focused either on pre-defined sets of genes (9–10) or on arbitrarily defined genomic intervals (8), the extent to which such relocalization occurs for TADs genome-wide remains unclear. An advantage of genome-wide chromatin conformation capture assays is the opportunity to define eu- and heterochromatic compartments in cell nuclei, and to analyze the dynamics of their relative arrangement (2, 8). To map the extent of such 3D reorganization genome-wide, we employed a previously established approach (2, 8) and assigned each domain to one of two major compartments, A and B, displaying eu- and heterochromatic features respectively (2). While compartmental assignment for most TADs was preserved following differentiation (**Fig. 1***C*–***D***, **Table S2, Fig. S8**), 302 (13.5%) domains switched compartment (**Fig. 1*D***), including those containing *Nanog*, a central pluripotency factor, and *Sox11*, a neural lineage marker. These domains moved from the A to B and from B to A compartments, respectively. We then investigated whether compartment changes were linked with the regulation of intra-TAD gene expression. We profiled ES and NS cells by RNA-seq and analyzed the alterations of TAD-wise expression levels. Generally, for TADs that switched compartment, the direction of transcriptional regulation matched the direction of compartmental change (**Fig. 1*E***, SI Methods). These results show a correlation between transcriptional regulation and compartmental association of TADs. However, only a small fraction of TADs altered compartmental membership, and many transcriptionally regulated genes were located in TADs that did not change. These results led us to explore other aspects of spatial reorganization that may be linked with gene expression.

### Alterations of the exposure of TADs to the inter-chromosomal space are widespread upon neural induction

Single-locus experiments have revealed preferential positioning of active genes towards the chromosome territory surface (25–26). Moreover, relocation of genomic loci with respect to the chromosomal surface has been observed for certain developmentally upregulated genes (12–14), suggesting a link between nuclear positioning and the control of transcription. However, a parsimonious view relating gene expression activity to location at chromosomal peripheral sites was challenged by the observation that transcriptional activity is not confined to the surface of chromosomes (27–28). These observations have raised the question of how often surface repositioning occurs genome-wide, and what general rules relate this phenomenon to gene activity. To measure exposure to inter-chromosomal space, we computed for each TAD the proportion of inter-chromosomal contacts among all of its long-range contacts (the inter-chromosomal contact probability index, ICP, 21). Comparing ICP in both ES and NS cell types, we detected significant changes for as many as 50% of TADs (moderated *t*-test, *q*-value < 0.1, **Fig. 2*A***, **Fig. S9**). Among those where ICP significantly increased was the *HoxD* locus (**Fig. S10**). This result is remarkable, given that relocation to inter-chromosomal space has been observed by FISH for only a fraction of alleles in the NS cell population (13), and it reflects the high sensitivity of comparative ICP analysis. We also detected significantly increased ICP for the TAD containing the *HoxB* locus, in line with its peripheral location in NS cells (14). Moreover, we observed higher ICP in ES than NS cells for 17 of the 20 centromeres included in the analysis (**Fig. 2*A***). Centromeres were previously shown to be preferentially located at nuclear-internal sites in human ES cells compared to the differentiated cells (29). These results suggest that repositioning of domains to the nuclear peripheral sites sequesters them from forming inter-chromosomal associations. To test this, we analyzed the relationship between ICP and alterations in Lamin B1 binding (11), which reflects dynamic tethering of TADs to the nuclear envelope. We observed significant anti-correlation between Lamin B1 binding and ICP (**Fig. 2*B***). This result demonstrates that comparative ICP analysis provides a sensitive measure of relocalization with regard to inter-chromosomal space. Moreover, the relationship between ICP and Lamin B1 binding indicates a depletion of inter-chromosomal contacts at nuclear peripheral sites.

**Fig. 2.**
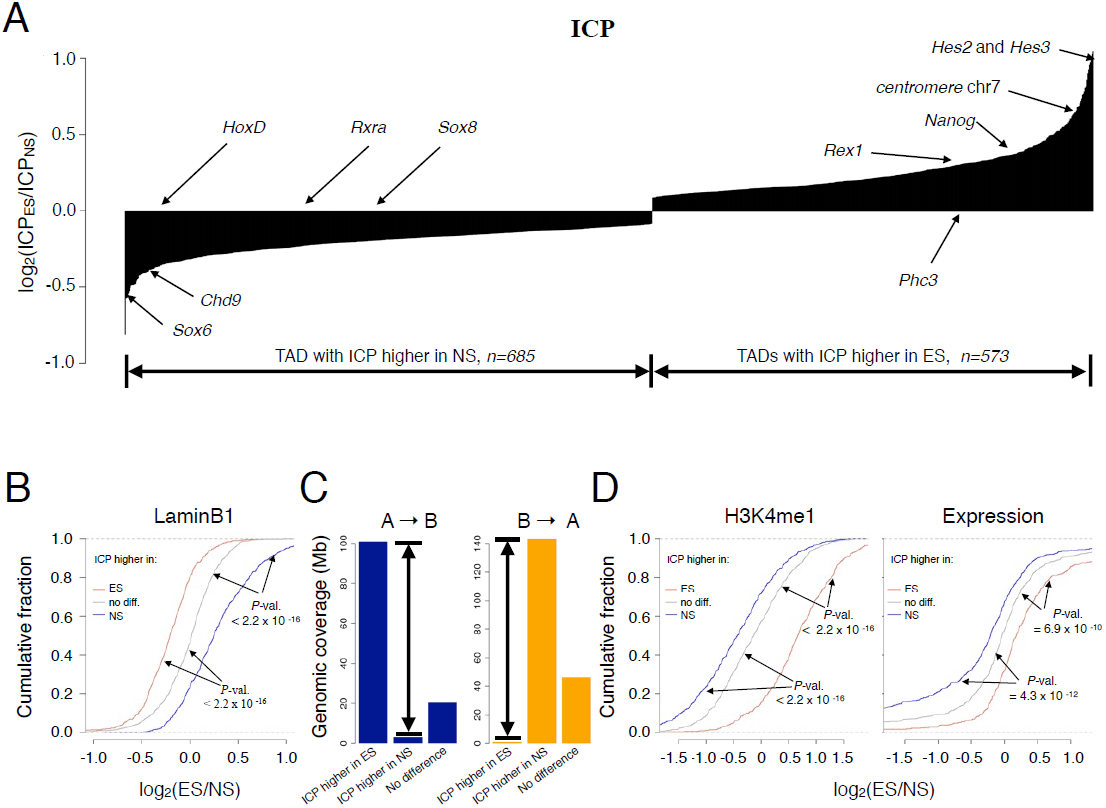
Relocation between chromosome-internal and exposed locations is widespread during differentiation. (***A***) Distribution of significant ICP changes of TADs. (***B***) Changes of ICP are anti-correlated with Lamin B1 binding. (***C***) TADs switching compartment display higher ICP in the condition in which they are in the active compartment (arrows). The *y*-axis depicts the cumulative genomic coverage of TADs that show compartment switching, bars are grouped by sign of ICP change. (***D***) Changes of ICP correlate with changes in H3K4me1 and expression.

### Internal organization and transcriptional activity within TADs correlates with exposure to inter-chromosomal space

Developmentally regulated gene looping outside chromosome territories has been associated with the activation of specific loci (12–14). We therefore investigated the genome-wide relationship between ICP changes and transcription. TADs with increasing ICP contained more up-than downregulated genes between the ES and NS cell states (Fisher’s exact test, *P*<10^-16^). Moreover, the subset of TADs that changed compartment membership displayed an activity-concordant shift in ICP (**Fig. 2*C***). In addition, alterations of TAD-wise H3K4me1 histone modification, a marker of active regulatory regions (30), as well as alterations in overall TAD expression levels correlated with ICP change (**Fig. 2*D***). Single-locus studies point to a correlation between locus activity and the formation of intra-chromosomal DNA loops (31–32). We therefore asked whether concurrent changes in domain activity and ICP could also be related to shifts in the internal organization of the domains. To address this question, we measured for each TAD the density ρ_int_ of strong and reproducible interactions within it (SI Methods). ρ_int_ was correlated with the average level of H3K4me1 per TAD (**Fig. 3*A***), reflecting a positive relationship between DNA loop formation and activity of cis-regulatory elements. Moreover, changes in ρ_int_ following differentiation were tightly associated with changes in ICP (**Fig. 3*B***–***D***).

**Fig. 3.**
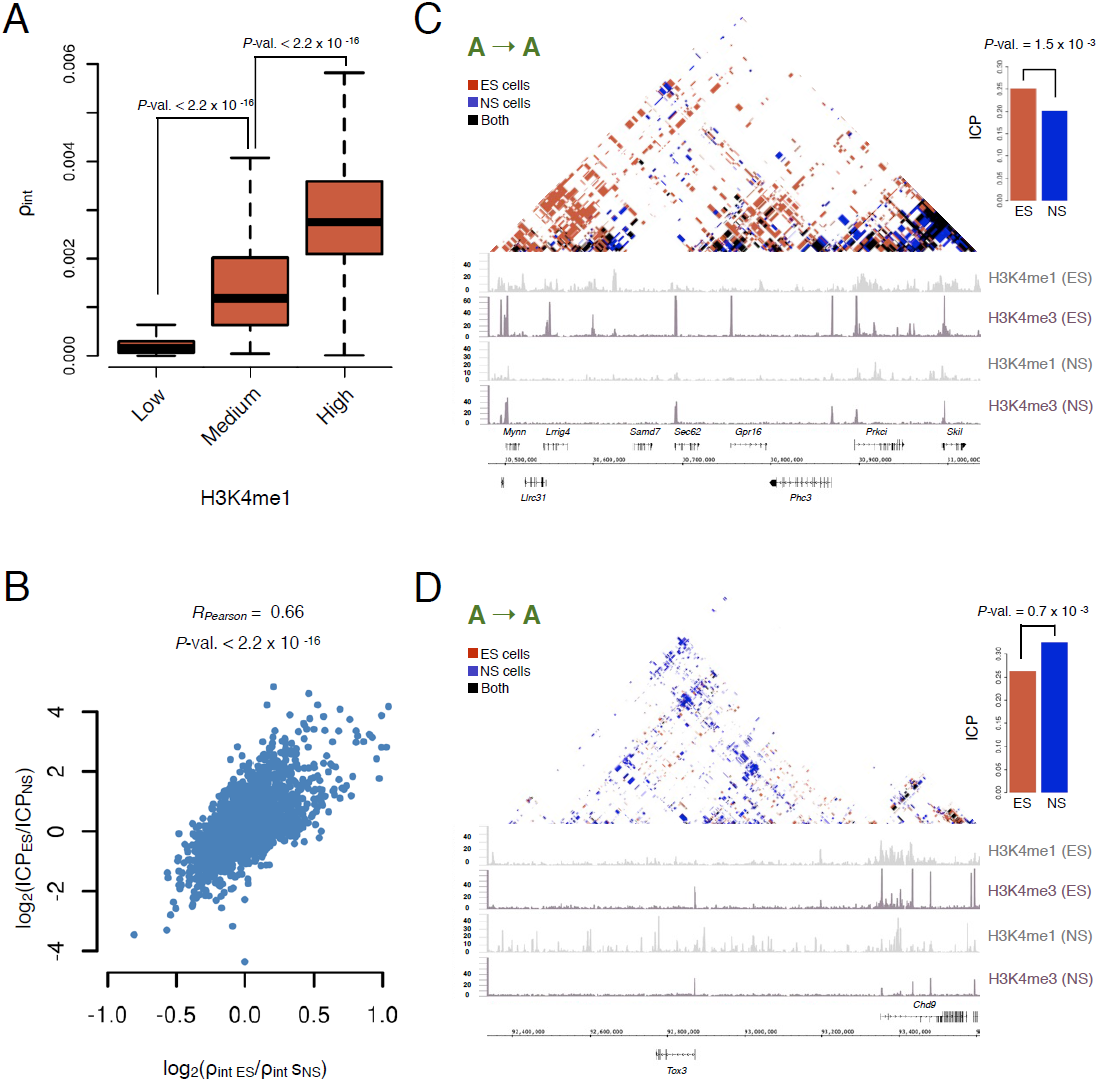
Changes in the internal architecture of TADs correlate with exposure to inter-chromosomal contacts. (***A***) TAD-wise average H3K4me1 levels correlate with ρ_int_, the density of strong and reproducible interactions (SI Methods). (***B***) Changes in ICP correlate with changes in ρ_int_. (***C***) Example of a TAD with significantly decreasing ICP, decreasing ρ_int_ and decreasing H3K4me1. (***D***) Example of a TAD with significantly increasing ICP, increasing ρ_int_ and increasing H3K4me1. Both examples (***C***-***D***) involve domains where compartmental membership was not altered. *P*-values: two-sided *t*-test.

Taken together, these results indicate that changes in the exposure of genomic regions to inter-chromosomal space are widespread in neural differentiation and coincide with changes in gene expression and internal TAD organization. Alterations of the ICP are not restricted to domains switching compartment: 52% of TADs remaining in the same compartment during neural induction exhibit altered ICP levels (**Fig. 2*A***, **Fig. 3*C***–***D***). Thus, changes in activity and strength of internal connectivity can be observed even for TADs where compartmental membership is preserved. The strong correlation between ICP and the number of interactions within TADs makes it tempting to speculate that the entropy of loop formation could mediate the relocalization of TADs towards the chromosomal periphery. However, the delineation of causal relationships between these processes remains an open question.

### Loss of pluripotency is associated with changes in chromatin organization

The regulatory mechanisms governing chromatin remodeling in pluripotent cells are crucial in the early development of multicellular organisms. Pluripotency has been associated with a specific chromatin state featuring higher overall activity and less condensed heterochromatin structure (33). On that basis, one might expect an increased size of the euchromatic A compartment in ES compared to NS cells; however we observed no such difference. The A compartment covered 905 and 957 Mb of the genome in ES and NS cells, respectively. We then sought to compare the inter-domain networks of intra-chromosomal interactions in ES and NS cells, to identify and characterize potentially unique features of chromatin organization in pluripotent cells. Quantitative comparison of chromatin interactome data is challenging. First, the data are affected by statistical sampling, and any observed difference must be weighted against the natural variability in these data. Second, the data can be affected by systematic measurement biases, such as: (*i*) the efficiency of the restriction enzyme digestion, (*ii*) CG bias and (*iii*) the cell cycle states of compared cell populations. All these factors affect the number of sequenced reads in a TCC experiment, and should be considered in order to obtain accurate estimates of quantitative differences in the inter-domain interactions.

We developed a statistical model that accounts for these factors (SI Methods, **Fig. 4*A***, 34). We detected 20,701 differential interactions (16% of all possible interactions), i.e. interactions between two TADs that significantly changed in strength between ES and NS cells (Wald-test, *q*-value<0.1). Nearly all TADs were involved in at least one differential interaction (**Fig. 4*A***–***B***, Fig. S11), underscoring the pronounced differences in global chromatin organization between ES and NS cells. Most of the differential inter-TAD contacts occurred between TADs from the same compartment (**Fig. 4*C***). For example, the TAD containing *Prdm14*, a pluripotency regulator (35), displayed significantly stronger interactions in ES cells, all of which associated it with the A compartment in ES cells. Interactions within the B compartment were more often stronger in the differentiated cell type, consistent with the less structured organization of ES cell heterochromatin (17, 33).

**Fig. 4.**
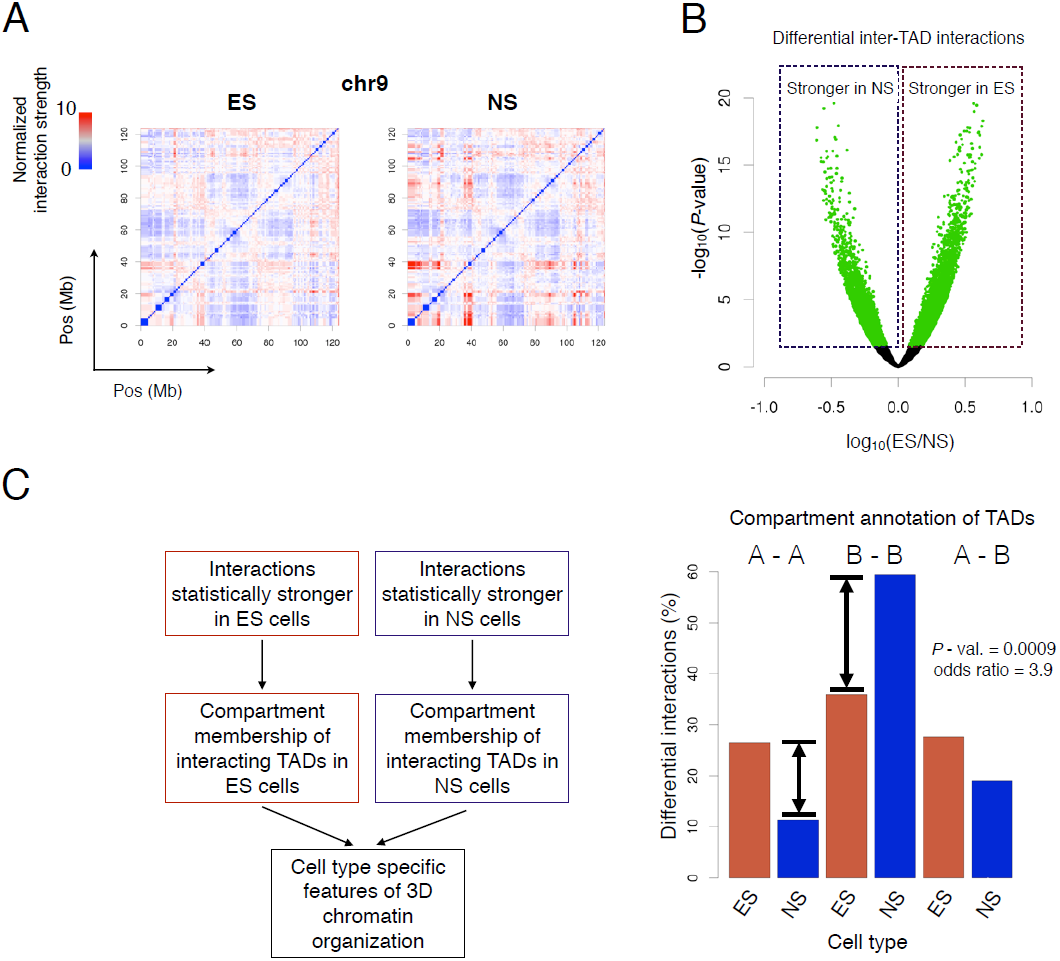
Comparative analysis of TAD interactomes reveals the more pronounced role of euchromatin in pluripotent nuclear organization. (***A***) Chromosome-wide normalized ligation frequency matrices at TAD resolution of chromosome 9 for ES and NS cells. Matrices of all chromosomes with two biological replicates per condition were used for comparative analysis. (***B***) Volcano plot of differential (ES vs. NS cells) intra-chromosomal inter-TAD interactions. (***C***) Categorization of the statistically significant differential inter-TAD interactions (A-A and B-B: both interacting TADs in the same compartment, A-B: interactions between compartments).

Surprisingly, interactions between TADs in the A compartment were twice as likely to be stronger in ES than in NS cells (**Fig. 4*C***), suggesting a different nature of the active compartment in pluripotent and differentiated cell types. We hypothesized that the increased strength of contacts between TADs from the A compartment might be related to higher activity of this euchromatic compartment in ES cells. Indeed, we found that TADs residing in the A compartment displayed higher levels of H3K4me1/me2/me3 in ES than in NS cells (**Fig. 5*A***), while no such trend existed for TADs remaining in the B compartment. Moreover, genes highly expressed in ES cells were more often found in the active compartment, compared to those highly expressed in NS cells (**Fig. 5*B***, Fisher’s exact test, P=3.4×10^-7^, **Fig. S12**). These results indicate that euchromatin in ES cells adopts a specific architecture whereby strong inter-TAD contacts are frequent. This may be a consequence of the higher overall activity of ES cell euchromatin and could play a role in maintaining the pluripotent state.

**Figure 5.**
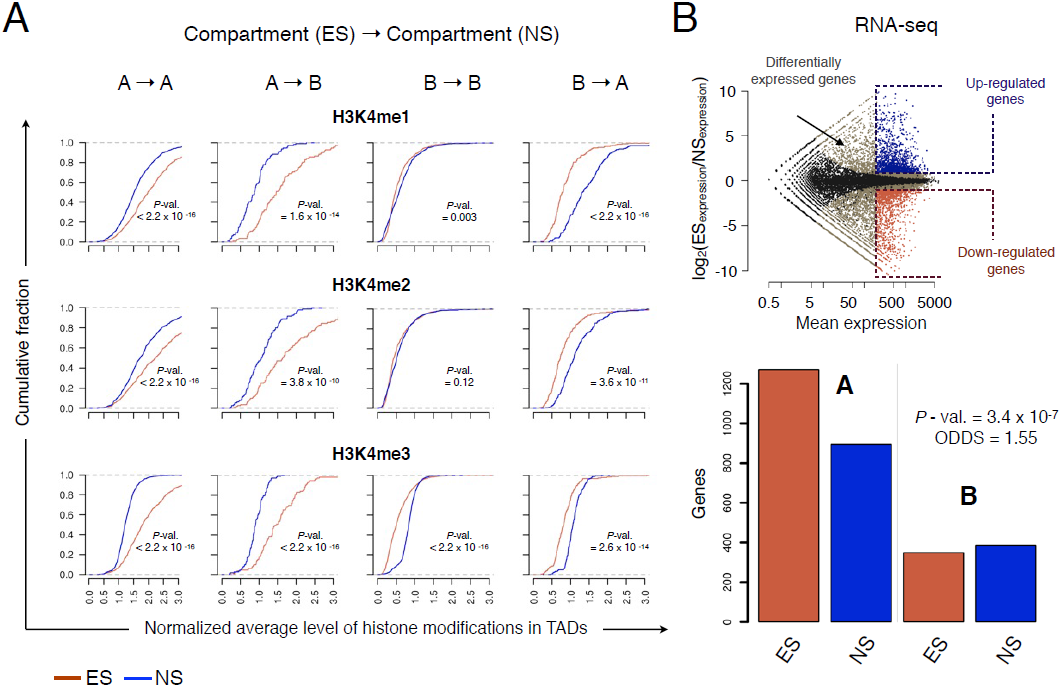
Euchromatin in ES cells displays higher overall activity relative to differentiated cells. (***A***) H3K4me1/me2/me3 histone modifications of euchromatin are stronger in ES cells than in NS cells (first column, A→A). The cumulative distribution functions show normalized average level of histone modifications per TAD. The other comparisons (B→B, A→B, B→A) are as expected, indicating adequate data normalization. *P*-values: two-sided *t*-test. (***B***) In ES cells, genes with higher expression are more often located within the euchromatic A compartment, compared to genes with higher expression in NS cells. Considered were genes with mean expression in the top 30%, fold change >2 or <1/2 and *q*-value < 0.1 (Wald-test).

### High-resolution analysis of chromatin interactome maps reveals frequent stability of promoter structure in cell differentiation

Promoters are central to the regulation of gene expression, receiving and consolidating the information provided by multiple *cis*-regulatory elements. Mapping of promoter interactomes is instrumental to dissect the mechanisms underlying complex transcriptional programs. Two recent studies considered promoter interactomes in different cellular contexts (16, 20). However these analyses focused only on associations involving Pol II-bound elements. A comprehensive assessment of 3D promoter interactions and dynamics in fundamental biological processes is still missing. To infer this from our TCC data, we identified genome-wide interactions that involved restriction fragments overlapping annotated promoter regions (± 5 kb around transcription start sites, **SI Methods**). We limited our analysis to non-overlapping promoters of protein coding loci. Surprisingly, despite the highly divergent cell identity and gene expression programs of ES and NS cells (**Fig. 5*B***), promoter interaction patterns were in many instances preserved. Prominent examples include *Nodal*, which promotes self-renewal in ES cells (**Fig. 6*A***, 36), and *Wnt7a*, essential for NS cell maintenance (**Fig. 6*B***, 37). Importantly, interactions between enhancers and promoters existed independent of whether those enhancers showed enrichment of H3K4me1 and H3K27ac marks, a signature associated with activity (**Fig. 6*A***–***B***, yellow zone 38–40).

**Figure 6.**
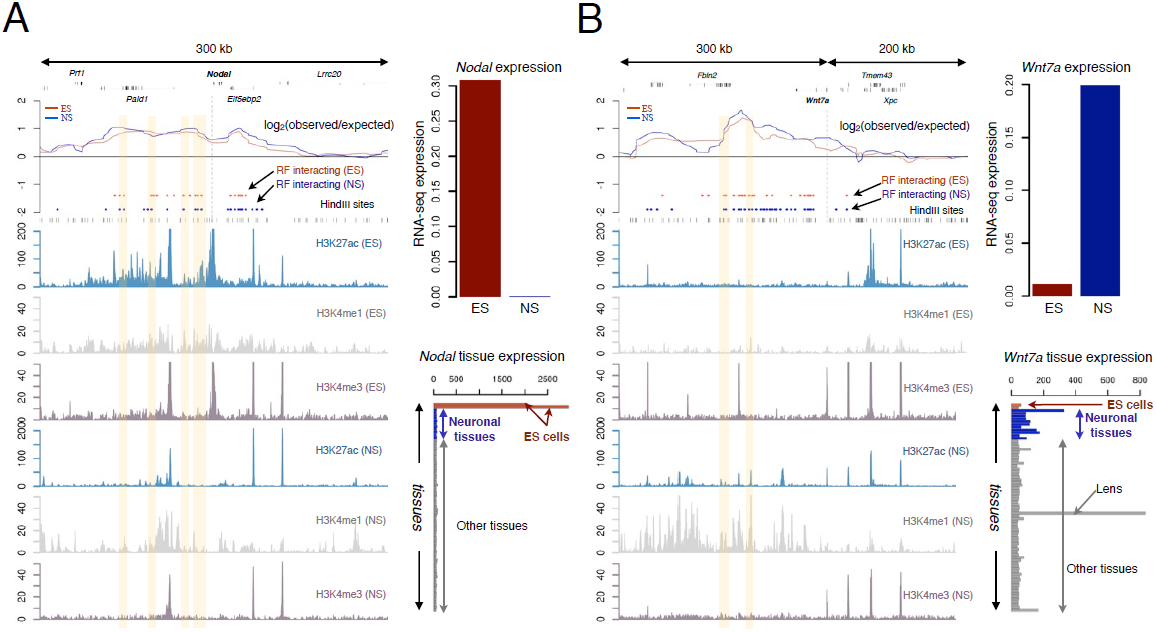
Promoter interactomes of transcriptionally regulated tissue-specific *Nodal* and *Wnt7a* genes. (***A***) TCC-based promoter interactome of *Nodal* in ES and NS cells (left panel). The profiles show the correspondence score between *Nodal* promoter and surrounding RFs normalized by genomic distance. Red and blue bars: RFs interacting with *Nodal* promoter in ES and NS cells respectively. Black bars: HindIII cutting sites. ChIP-seq tracks are from refs. 39, 64. Yellow: activity-regulated enhancers stably interacting with the promoter. Upper right panel: expression of *Nodal* in ES and NS cells. Lower right panel: expression of *Nodal* in mouse tissues. (***B***) Promoter interactome of *Wnt7a*; annotation as in ***A***.

### Widespread maintenance of promoter-enhancer interactions

Enhancer-promoter communication is a key determinant of cell identity (1, 7, 15, 38). We therefore investigated the extent of structural preservation in promoter-enhancer interactions (PEI) in the context of differentiation. We considered a set of 51,964 restriction fragments overlapping enhancers that were identified in ES and NS cells (39) and inferred, from our TCC data, 42,042 PEI with 12,861 promoter regions corresponding to 8,325 genes (**Fig. S13**, SI Methods). Multiple lines of evidence pointed to a high accuracy of our PEI network: (*i*) 98% of PEI occurred within TAD boundaries (7), (*ii*) we recovered the reported PEI for the *Oct4* (41), *Phc1* (42) and *Nanog* (43) genes in ES cells (**Fig. S14**), (*iii*) we correctly detected the enhancer-switching event for the *Sox2* gene (**Fig. S15**, 20), (*iv*) genes with PEI were expressed at higher levels than genes without a detected PEI (**Fig. S16**), (*v*) we observed an enrichment of functional annotations related to pluripotency for genes contacted by enhancers active in ES cells, and to neurogenesis for those active in NS cells (**Figs. S17 and S18**).

To investigate the relationship between contact formation and enhancer usage, we focused on activity-regulated enhancers (39) and divided these elements according to chromatin dynamics (**Fig. 7*A***). Instances of activity-independent PEI were frequently observed. For 10886 (∼40%) of PEI involving regulated enhancers, proximity was detected in both cell types, including in the condition where, according to histone modification status, the interacting enhancer was inactive (**Fig. 7*A***). The same proportion of stable PEI was observed within each group of enhancers (**Fig. 7*A***), indicating that structural maintenance of PEI is equally frequent in induced and repressed distal elements.

**Figure 7.**
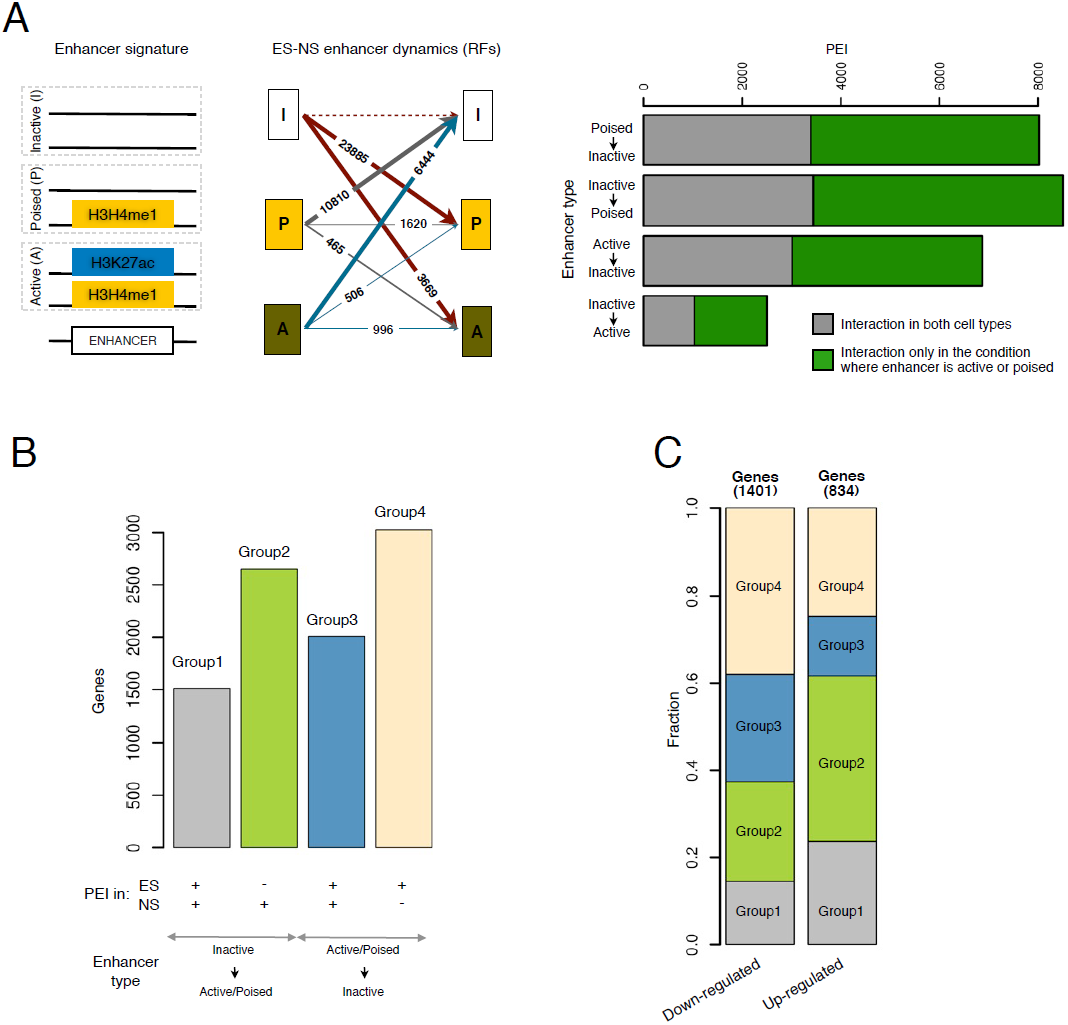
Many promoter-enhancer interactions (PEI) exist in an enhancer-activity-independent manner. (***A***) Left: definition of enhancer signatures. Middle: changes in histone modifications at enhancers during the ES-to-NS cell transition. Thick arrows: enhancer-containing RFs used for further analysis. Right: PEI stratified by enhancer chromatin signature dynamics. (***B***) Gene-level PEI dynamics involving activity-regulated enhancers. Per gene, PEI for all annotated promoters were considered. (***C***) PEI dynamics involving activity-regulated enhancers for differentially expressed genes. Down- and upregulated genes were stratified according to whether any stable PEI with an activity-regulated enhancer was detected.

To gain more insight into the involvement of stable PEI in the regulation of gene expression, we classified genes based on whether a preserved interaction was detected at those loci and how the activities of these enhancers changed (**Fig. 7*B***). This revealed roughly equal sized groups of genes with and without structural preservation of PEI following neural differentiation. For instance, *Fgfrl1* (Group 1) displayed in both cell types a PEI with an enhancer active in NS cells alone, whereas the PEI of *Oct4* (Group 4) with an ES cell-specific enhancer was present only in ES cells and the PEI of *Pax6* (Group 2) with an NS cell-specific enhancer was exclusive to NS cells (**Figs. S19** and **S21**). Loci with preserved PEI were associated with functions in early development (BH-adjusted *P*<0.01, **Fig. S22**), suggesting that structural maintenance plays a previously unappreciated role in the developmental regulation of a large fraction of the genome. Indeed, 1016 (33%) of genes with stable PEI were differentially expressed (*q*-value<0.1, FC>2) (**Fig. 7*C***, Groups 1 and 3 for down- and upregulated genes respectively). Considering all differentially expressed genes, we observed that among down-regulated genes, those contacted by repressed enhancers (Group 3 and 4) were overrepresented, while among up-regulated genes loci contacted by induced enhancers (Group 1 and 2) were more prevalent (**Fig. 7*C***). Thus, interaction with an active or poised enhancer is predictive of transcriptional status (**Fig. S16**). However, the silencing of distal regulatory elements often does not result in a reduction of promoter interaction strength.

For a subset of genes, the cell type in which a PEI was observed was inconsistent with both the direction of transcriptional regulation and enhancer activity change (**Fig. 7*C***, Groups 2 and 4 for down- and upregulated genes, respectively). However, the promoters of these loci contacted greater numbers of active or poised enhancers (SI Methods) in the condition were the gene was expressed at a higher level (*P*-value = 8.3 x 10^-13^ for genes in Group 2 and *P*-value = 3.9 x 10^-12^ for genes in Group 4 of **Fig. 7*C***, two-sided *t*-test). This result suggests that in these cases, the PEI imparted a minor influence on gene expression, or that the PEI in the contrasting cell type did not reach the detection threshold.

### Stable PEI are preferentially detected at complex, multi-enhancer loci

To further assess the functional implications of PEI stability in the regulation of gene expression, we analyzed the chromatin signature of promoters based on a detected PEI involving an activity-regulated enhancer. We did not detect a significant difference in the levels of H3K27ac, a modification characteristic of active promoters (44–45), or in Pol II enrichment (**Fig. 8*A***–***B***, SI Methods, see as well **Fig. 6*A***–***B***). Spatial proximity of NS-specific enhancer elements already present in ES cells could convey an advantage in the efficiency of transcriptional upregulation of the *cis*-linked neural genes. This would be reflected by a more rapid onset of gene expression during stepwise neural fate induction. We analyzed published data reporting transcriptional changes during neuronal differentiation of ES cells (46). However, we did not detect a difference in the kinetics of up-regulation of gene expression between genes with stable PEI involving activity induced enhancers compared to genes that featured only cell NS cell-specific PEI.

**Figure 8.**
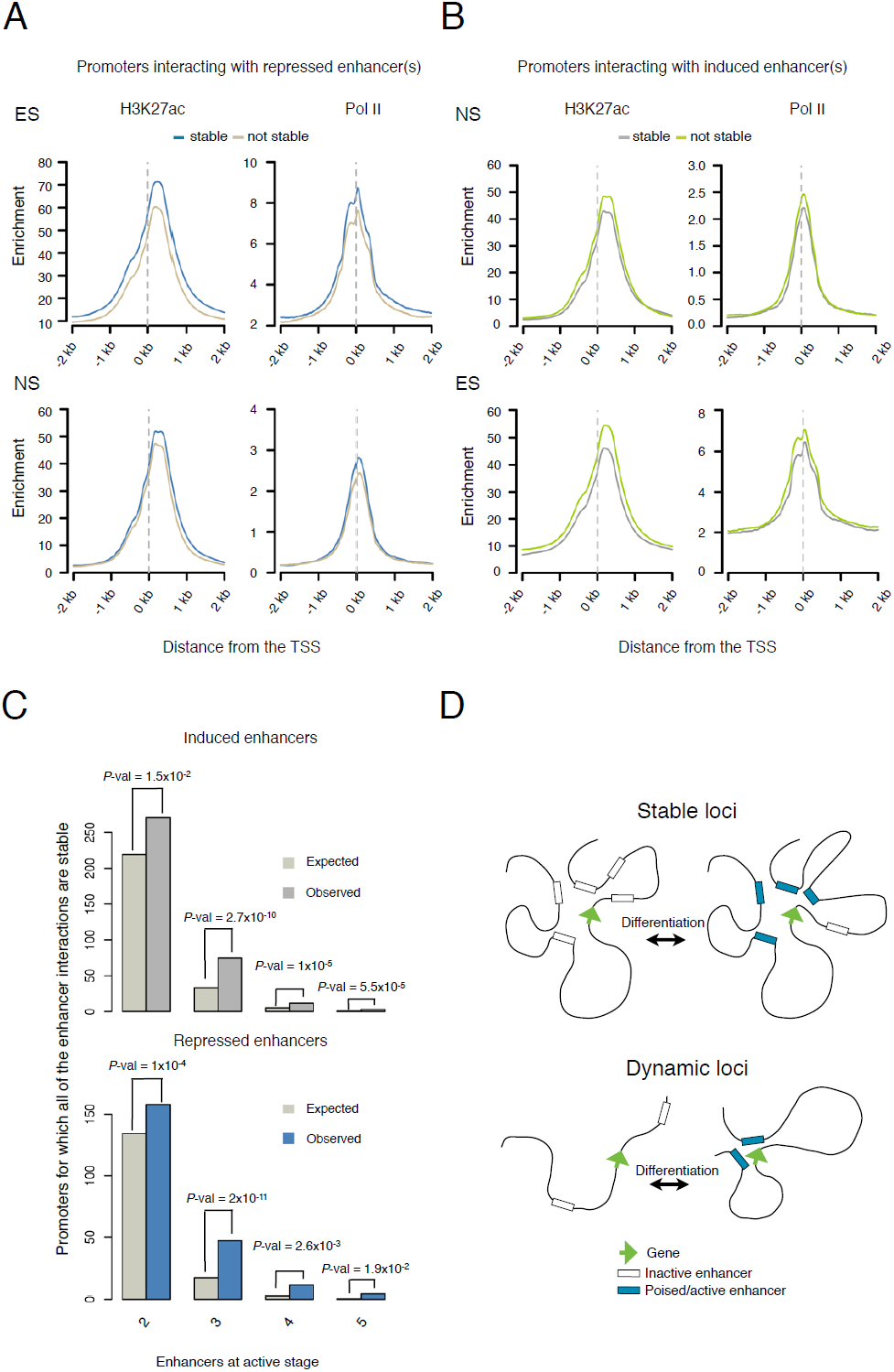
PEI stability is pronounced at complex, multi-enhancer loci. (***A***) Average profiles of H3K27ac marks based on ChIP-seq data and Pol II at promoters involved or not in stable PEI with repressed enhancers and in (***B***) for promoters involved or not in a stable PEI with induced enhancers. (***C***) Loci interacting with multiple enhancers show an overrepresentation of exclusively stable PEI. Bar plots are stratified by number of interacting active enhancers and cell type in which they are active. *P*-value: *Chi*-Square test. (***D***) Model depicting the role of the stability of promoter-enhancer interactions in transcriptional regulation. Complex loci interacting with multiple enhancers more often display PEI stability than those contacting few enhancers.

CTCF and cohesins have been implicated in the establishment of cell type-invariant interactions (18). To test whether CTCF is involved in the maintenance of stable PEI, we isolated CTCF peaks (47) in ES and NS cells and focused on the constitutive CTCF-enriched regions. We divided the induced and repressed enhancers for which we observed a PEI into two groups defined by whether or not the PEI was stable before and after neural differentiation. We found no difference between the two groups. In both cases, around 18% of enhancers were overlapping constitutive CTCF binding sites.

The presence of stable promoter interactions may arise due to more complex, intrinsic differences in the structure of the *cis*-regulatory network. We hypothesized that structural stability may be related to the complexity of the promoter interactome of a given gene. Indeed, we found that multi-enhancer loci were enriched for stable PEI (*P*-value = 5.3 x 10^-16^ and *P*-value = 1.7 x 10^-14^ for promoters contacting induced and repressed enhancers, respectively; Fisher’s exact test, SI Methods, **Fig. 8*C***). In summary, these results show that dissociation between enhancer activity and the formation of physical contact with promoters is widespread in key developmental processes such as neural lineage commitment and differentiation. We suggest that preserved chromatin architecture serves to ensure efficient regulation of a complex *cis*-regulatory landscape (**Fig. 8*D***).

## DISCUSSION

Here we reveal the dynamic alterations of chromatin conformation at multiple genomic-length scales in an *in vitro* model of neural cell fate specification. On a global scale, we show that repositioning of genomic domains with respect to chromosome surfaces is prevalent and associated with transcriptional regulation. We demonstrate for the first time that such widespread alterations of chromatin structure are initiated in mammalian cell differentiation, and show direct evidence of a genome-wide link between the positioning of domains at chromosomal surfaces and transcriptional activity.

The mechanisms underlying movements of active TADs to the surface of the chromosome are largely elusive. We suggest that DNA secondary structures produce entropic effects that drive repositioning away from the chromosomal interior. Indeed, transcription factors and Pol II have access to the entire chromosome territory (48). Moreover, as recently shown, reprogramming-mediated induction of *Nanog* and *Lefty1* expression is preceded by the formation of DNA loops connecting promoter and enhancer elements (61). Similar long-range structures could thus favor exposure of those regions to the chromosomal surface where additional events, initially independent of expression, occur. For instance, pluripotency factors such as Klf4 have been implicated in the maintenance of inter-chromosomal interactions in the context of reprogramming (60). Interestingly, Klf4 binding and the onset of inter-chromosomal interactions in ES cells precede transcriptional up-regulation of the target *Oct4* gene. Taken together, these observations suggest a model whereby the alteration of 3D chromatin structure is initiated by an increase in the activity of regulatory elements and precedes transcriptional activation. These alterations include both internal reorganization of TAD structure and its exposure to inter-chromosomal space.

Our data indicate an overall tendency of active genomic regions to be exposed to inter-chromosomal contacts. This view agrees with the recently published observation of frequent inter-chromosomal interactions between active enhancers and promoters (20). The presence of transcription factories (49), i.e. regions of active transcription, at sites of inter-chromosomal interactions (50–51) could provide additional stabilization forces that maintain the peripheral localization of active TADs. The widespread exposure of actively transcribed genes could globally reinforce co-regulatory programs of gene expression. Further studies are required to delineate the causal relationship between these features.

Chromatin organization of pluripotent cells has been studied extensively (33, 52). Analyses of the mobility of chromatin proteins revealed low retention time in ES cells in comparison to differentiated cells (53), and the hyper-dynamic character of binding of structural proteins to chromatin in ES cells has been suggested to play a functional role in the maintenance of the pluripotent state (53). Heterochromatin in particular has been shown to be less condensed in ES cells (52–53), and chromatin interaction patterns obtained by Hi-C and 4C-seq revealed reduced formation of long-range contacts of heterochromatic regions (17). By analyzing intra-chromosomal inter-TAD interaction strengths in ES and NS cells, we confirm the latter result and discover a new feature of chromatin organization: pronounced, activity-linked contacts within euchromatin in pluripotent cells. Based on a comparison of the enrichment of active histone modifications (30) between ES and NS cells, along with transcriptional activity, we suggest a link between this unusual chromatin arrangement and increased gene expression. Widespread transcriptional activity in ES cells has been shown to include lineage-specific factors and normally silenced repetitive elements (54). These are expressed at low levels (54) and are usually located within the heterochromatic compartment B. Here we show an ES cell-specific bias of highly expressed genes towards the euchromatic compartment A. We suggest that reinforcement by spatial clustering of euchromatin is a specific property of ES cells. These findings may contribute to a better understanding of the mechanisms promoting pluripotency.

In contrast to proposed models of promoter-enhancer interactions which stipulate activity-dependent formation of PEI (15, 31–32), our high-resolution analysis of chromatin interaction patterns reveals widespread dissociation between enhancer element activity and formation of contacts with promoters. Studies of several developmentally regulated genes showed that PEI can form in the absence of apparent enhancer activity (55–56). Moreover, such architectural preservation was recently reported in response to environmental cues (57–59). Our analysis supports the hypothesis of the permissive model for the modulation of gene expression by higher-order regulatory interactions (56). We show that dissociation between PEI and enhancer activity is widespread in lineage commitment and cell differentiation. Preservation of contacts between *cis*-regulatory elements thus appears to be a fundamental determinant of these dynamic cellular processes. Here, we show that ∼40% of PEI are maintained despite regulation of enhancer activity. Previous analyses of chromatin interactome maps in ES and NS cells at individual loci (18) have revealed that cell-type invariant interactions can be mediated by CTCF/cohesin complexes (18). However, our genome-wide analysis reveals that maintenance of CTCF binding appears to explain only 18% of stable PEI events. The mechanisms underlying the remaining set of stable PEI remain elusive and require further characterization. One possibility is that the overall structure of a TAD, i.e., the spatial arrangement of adjacent interactions, tethers regions together and can lead to stable PEI formation. An alternative explanation implies the action of yet un-identified transcription factors in the maintenance of stable PEI.

CTCF is thought to play a fundamental role in 3D chromatin organization (62–63); in this context, our result showing that constitutive CTCF binding is similar for stable and dynamic PEI is surprising. Recent studies of somatic cell reprogramming show that the establishment of pluripotent specific long-range interactions precedes the transcriptional upregulation of key pluripotency factors (60–61). It is thus conceivable that CTCF is involved in the dynamics of PEI formation, for instance by ‘priming’ certain interactions to be formed early (e.g. at the neuro-ectodermal cell fate commitment step) during differentiation. Kinetic measurements of genome-wide PEI during cell fate transitions and functional experiments on the role of CTCF in the establishment and maintenance of PEI would be instrumental for a deeper understanding of these processes.

## Acknowledgements

We thank V. Benes and B. Haase of the EMBL Genomics Core Facility, A. Riddell and A. P. Gonzalez from the EMBL Flow Cytometry Core Facility, A. Reyes for help with data handling, F. Spitz for critical reading of the manuscript. We thank S. Pollard for helpful advices.

## Author Contributions

A.P. performed all experiments, A. P. and W. H. designed the study with input from P.B. and L.M.S. A. P., B. K. and W. H. performed the data analysis, F. A. K. and M. O. performed the initial data processing. P. B. contributed to the experiments, L. M. S. and S. A. contributed to the data analysis, A. P. and W. H. wrote the manuscript with input from the other authors.

## Methods

**Cell culture**. 46C cells were cultured in Glasgow Modified Eagles Medium (GMEM, Invitrogen), supplemented with 10% (v/v) fetal bovine serum (FBS) (Sigma), 2 ng/ml of inhouse produced leukaemia inhibitory factor (LIF) and other nutrients, on gelatin-coated (0.1% (v/v)) dishes. Differentiation to neural stem cells was performed by plating ES cells onto gelatin-coated dishes and incubating with N2B27 medium in the absence of LIF. After five days, GFP+ cells (the *Sox1* expressing cells) were sorted by flow cytometry and seeded onto laminin coated dishes. Cells were expanded in N2B27 medium supplemented with recombinant EGF and bFGF (final concentration of 10 ng/ml) until loss of GFP expression and uniform gain of Nestin expression was observed. Detailed ES and NS cell culture protocols are provided in the Supplementary Information.

**Tethered Chromatin Conformation** was performed as described previously (21). Paired-end sequencing was performed with Illumina HiSeq2000.

**RNA-seq**. Total RNA was isolated using Trizol (Invitrogen) according to manufacturer’s recommendations. Traces of DNA were removed by Turbo DNaseI enzyme (Ambion) treatment. Strand-specific RNA-seq libraries were prepared with the TruSeq (Illumina) kit and sequenced with Illumina HiSeq2000.

**Computational methods**. Analytical procedures are detailed in the Supplementary Information, along with a complete transcript of the analysis in the form of a documented R script (‘vignette’). The vignette source files and all required input data are provided in the experiment data package *ESNSTCC* on www.bioconductor.org.

